# Viability and mitochondrial bioenergetic functions in human colon cancer cells are not affected by treatment with peptides mtCPP1, UPF25, mtgCPP, and mtCPP1gHO

**DOI:** 10.1101/2020.07.03.185967

**Authors:** Natalja Timohhina, Carmine Pasquale Cerrato, Kaido Kurrikoff, Vladimir Chekulayev, Jekaterina Aid-Vanakova, Marju Puurand, Kersti Tepp, Igor Shevchuk, Ulo Langel, Tuuli Kaambre

## Abstract

The popularity of the specially synthesized cell-penetrating peptides (CPPs) in cancer treatment has grown recently. The main aim of this study was to investigate the effects of four mitochondrially targeted antioxidant CPPs on the viability and bioenergetic function of mitochondria in human adenocarcinoma Caco-2 cells. The number of viable cells was measured by MTT and trypan blue assays. Respirometry and the permeabilized cell technique were applied to measure the mitochondrial function in this cell line. We did not observe any significant effect of CPPs on the mitochondrial reserve respiratory capacity, the function of respiratory chain complexes, and the inclination of these cancer cells to aerobic glycolysis. mtgCPP peptide with the highest antioxidant activity demonstrated improved mitochondrial coupling efficiency. CPPs do not affect mitochondrial function directly but can be considered in therapeutics as a drug-delivery molecule

## 1. Introduction

Malignant transformation of cells is associated with thorough reprogramming in the metabolic pathways involved in the production of high-energy compounds, as well as in the biosynthesis of cellular lipids, proteins, and nucleic acids [1-3]. Unlike normal cells, cancer cells may metabolize glucose to lactate at high rates even in the presence of oxygen, defined as the Warburg effect [4, 5]. Not all cancer cells rely on glycolysis as their major source of ATP and, because of the oxidative phosphorylation (OXPHOS) system is essential for tumor growth [6-10].

Targeting tumor metabolism is a challenging diagnostic and therapeutic strategy for imaging and selective elimination of cancer cells [8, 11]. Mitochondria are the promising objects for that due to their specific role in cancer energy metabolism. The targets for the mitochondrial treatment are hexokinase (HK), thiol redox status, VDAC (voltage-dependent anion channel), adenine nucleotide translocase, electron transport chain, mitochondrial inner membrane and tricarboxylic acid cycle [12]. A big challenge is to overcome not so easily penetrable cellular membrane to excess intracellular compartments. Therefore, there is an urgent need for a precise drug-delivery system into mitochondria and the requirement of such a transport mechanism that does not affect the mitochondrial metabolism in an unknown way.

Cell-penetrating peptides (CPPs) are peptides with a sequence length usually between 4 and 30 amino acids. They possess cationic and amphipathic properties, needed characteristics to facilitate the delivery of the cargo molecule. Due to their low cytotoxicity and their possibility to transport many different types of bioactive cargo – such as peptides, proteins, drugs liposomes, phages, plasmid, nucleic acids - they have emerged as a new approach to the treatment of various infection and non-infectious diseases, including cancer [14-18]. Some CPPs can exert a strong antitumor effect or serve as the carriers of polar cytotoxic agents inside malignant cells [14, 19, 20].

Recently, we have synthesized series of mitochondria-targeted (mtCPP), glutathione analog peptides (UPF), and dual antioxidant peptide mtgCPP by fusion of mtCPP1 and UPF25 [21-23] to protect these organelles from oxidative damage [23, 24]. We have shown that mtCPPs have a superoxide anion scavenging ability in different cell lines, such as HeLa 705 (human cervical carcinoma), bEnd.3 (mouse brain endothelial), U87 (human primary glioblastoma), CHO (Chinese hamster ovary). Even stronger antioxidant activity was detected of the fused peptide mtgCPP compared to mtCPP1 and UPF25 alone [23]. These series of mitochondria targeting peptides also exerted a protective effect against exogenously added hydrogen peroxide. The viability of any of the cell lines used in the studies was negatively affected by the treatment with any of these peptides. Colon adenocarcinoma remains the leading cause of morbidity and mortality worldwide. Unlike other types of cancer, colorectal cancer (CRC) partly relies on the mitochondrial OXPHOS as an energy source. Our previous studies showed that CRC cells, unlike many other human cancers, have high rates of basal and ADP-stimulated respiration along with obvious signs of stimulated mitochondrial biogenesis [9, 10]. Thus, controlling the mitochondrial respiration may be a new strategy for the treatment of CRC [25]. Moreover, it was reported that the intracellular level of ATP is a crucial factor determining chemoresistance in CRC [26]. The influence of the studied CCPs on the mitochondrial function of CPC has not been previously reported. These peptides could suppress the growth of colorectal carcinomas by inhibiting their mitochondrial respiration and/or production of reactive oxygen species (ROS). It is also important to note that recently some specially synthesized peptides have been proposed for the diagnosis of CRC and its treatment as direct cytotoxic agents or carrier molecules [27].

Our study aimed to test the effect of four CPPs with strong antioxidant abilities (mtCPP1, UPF25, mtgCPP, mtCPP1gHO) [23] on the mitochondrial bioenergetic parameters in human CRC cells (Caco-2 line).and to determine the influence of the peptide on the cell viability.

## 2. Materials and methods

### 2.1. Chemicals

Eagle’s Minimum Essential Medium (MEM) containing 1.5 g/L sodium bicarbonate, non-essential amino acids, L-glutamine, and sodium pyruvate was purchased from Corning (REF 10-009-CVR). Trypsin-EDTA solution, heat-inactivated fetal bovine serum, and antibiotics (penicillin, streptomycin, and gentamicin) were obtained from Gibco Life Technologies (Grand Island, New York, USA). Unless otherwise indicated, all other chemicals were purchased from Sigma-Aldrich.

### 2.2. Peptide synthesis, purification, and analysis

Solid-phase peptide synthesis, purification, and analysis were performed as described previously [23]. Briefly, peptides were synthesized using standard Fmoc chemistry on microwave-assisted synthesizer (Biotage, Sweden). The peptides were cleaved from the resin, precipitated in ether, and lyophilized. CPPs were also labeled with 5-(6)-carboxyfluorescein (FAM) to obtain FAM-mtCPP1, FAM-UPF25, FAM-mtgCPP, and FAM-mtCPP1gHO (molecular weight: 983.97, 842.5, 1449.5, and 2420.3, respectively) for microscopy studies. The peptides were purified using RP-HPLC on a preparative C8 column with H_2_O-acetonitrile (both solution containing 0.1% TFA) gradient. The identity of the final products was confirmed using UHPLC-MS (Agilent 1260 Infinity, Agilent Technologies, Santa Clara, California, USA). Stock solutions of the studied peptides (at final concentrations of 0.1 and 1 mM) were prepared in sterile mQ-water and stored at -20 °C.

### 2.3 Cells

Caco-2 cells were obtained from the American Type Culture Collection (HTB37™). These cells were grown in T75 flasks (Greiner Bio-One) in MEM supplemented with 10% FBS, 100 units/ml penicillin, 100 μg/ml streptomycin, and 50 μg/ml gentamicin at 37 °C in an incubator with 5% CO_2_ in the air. Cells were harvested by mild trypsinization followed by centrifugation at 150g for 7 min. The cell pellet was suspended in medium-B [25] supplemented with 2 mg/mL fatty-acid free bovine serum albumin and 5 μM leupeptin (a protease inhibitor) and stored on ice. The viability of these cells was around 95% according to trypan blue (TB) exclusion test.

Cells were treated with CPPs at 5 μM concentrations for 24 hrs at 37 °C, and 5% CO_2_ in the air before every assay.

### 2.4. Viability assays

Along with the TB test, cell viability was estimated by MTT Cell Proliferation Assay Kit (ATCC® 30-1010 K). Caco-2 cells were seeded into Falcon® 96-well microplates (from Corning) with a flat bottom at a density of 5000 cells/well and cultured at 37 °C with 5% CO_2_ for 24h. After that growth medium was replaced with new media containing different CPPs, and cells were incubated overnight. Under the same conditions, growth medium supplemented with sterile milli-Q water was used as a control. The optical density in each well was measured at 570 nm using 630 nm as reference wavelength on a FLUOstar Omega (BMG LABTECH, Germany) microplate reader.

### 2.5. Assessment of the mitochondrial respiration in Caco-2 cells

The mitochondrial respiratory activity in Caco-2 cells was evaluated using the saponin-permeabilized cell technique [28].

O_2_ consumption rate was measured at 25 °C using a high-resolution Oxygraph-2K respirometer (Oroboros Instruments, Austria); the solubility of oxygen was taken as 240 nmol/mL [29]. Respiration was measured as a specific oxygen flux (in nmoles O_2_/min/million cells) and normalized per mg of cellular protein. The protein concentration in cell lysates was determined using the Pierce BCA Protein Kit. The coupling of hexokinase (HK)-catalyzed processes with the OXPHOS system in permeabilized Caco-2 cells was assayed by oxygraphy via activation of mitochondrial respiration by locally generated ADP [10, 30]. This effect of glucose on mitochondrial respiration was expressed by glucose index (I_GLU_) that was calculated according to the equation (1): 

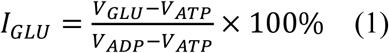

where V_ADP_ is the rate of O_2_ consumption in the presence of 2 mM ADP, V_GLU_ is respiration rate with 10 mM glucose, and V_ATP_ is respiration rate upon addition of 0.1 mM ATP. This index indicates the degree to which glucose-mediated stimulation of mitochondrial respiration is comparable to maximal ADP-activated rates of O_2_ consumption.

The coupling of adenylate (AK)-catalyzed processes with OXPHOS was evaluated by respirometry as described previously [10, 30]. AK index (I_AK_) determines the magnitude of this efficiency and is calculated according to the equation (2): 

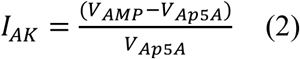

where V_AMP_ and V_Ap5A_ are rates of O_2_ consumption in the presence of 2 mM AMP and 0.2 mM diadenosine pentaphosphate (Ap5A, an inhibitor of AK), respectively.

### 2.6. Statistical Analysis

All data are expressed as mean values ± standard error of the mean (SEM) of at least 4-5 independent experiments. Statistical significance was assessed by one-way ANOVA followed by Holm-Sidak *post-hoc* test using SigmaPlot 11.0 software.

## 3. Results and discussion

### 3.1. Toxicity of CPPs on Caco-2 cells

To evaluate the cytotoxic activity of four different CPPs against human epithelial colorectal adenocarcinoma cells (Caco-2) were incubated with a dose of 5μM for 24h. Cell viability was determined by the TB exclusion test and MTT assay. Studied FAM-labeled CPPs did not affect the Caco-2 cells proliferation rate no difference was observed between control and CPP-treated cells (**Fig. 1)**. Similar effects were observed with HeLa 705, U87, and bEnd.3 cell lines as reported earlier [23, 31].

**Figure 1.**
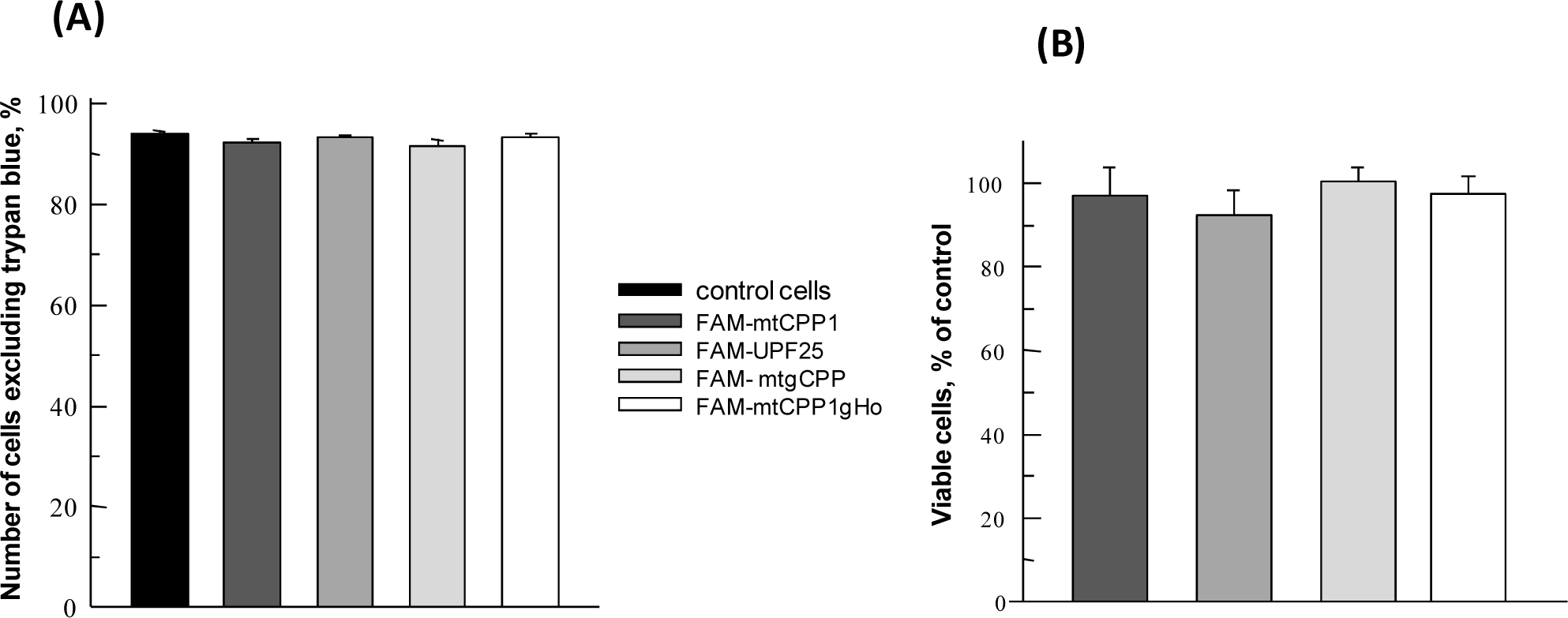
The effect of cell-penetrating FAM-labeled peptides at final concentrations of 5 μM on the viability of Caco-2 cells in trypan blue exclusion assay (**A**) and their proliferative activity that was estimated by MTT test (**B**). Cells were treated with the indicated peptides in complete growth medium for 24 hrs at 37 °C. Data are shown as mean values ± SEM, n = 4.

### 3.2. Effect of CPPs on the coupling of OXPHOS with hexokinase- and adenylate kinase-catalyzed processes in Caco-2 cells

Human CRC cells compared with unaffected tissue cells have an increased energy demand, which is manifested in their increased glycolytic activity and rates of OXPHOS [9, 25]. Colorectal adenocarcinoma maintains high rates of glucose consumption and has a higher tendency to aerobic glycolysis [9, 32]. This phenomenon may be related to the elevated expression of some glucose transporters [33] and glycolytic enzymes [34, 35], or predominantly, by the mitochondrial VDAC-bound HK-1 or 2, which use mitochondrially generated ATP to phosphorylate glucose in large amounts [5, 9, 10]. In this work, we evaluated Caco-2 cells as a model system expressing VDAC-associated HK-2 similarly to human colorectal cancer [25]. Experiments were carried out to estimate the impact of CPPs on the inclination of Caco-2 cells to aerobic glycolysis (the Warburg behavior). The addition of 10 mM glucose to permeabilized cells caused an increase in the respiration rate in both the CPP-treated and control cells but there is no significant difference between those groups (**Fig. 2**). Thus, the close coupling between the HK and OXPHOS system was maintained in all experiments. The effect of glucose consumption was expressed by the glucose index that reflects the degree of glucose response relative to the maximal rate of ADP-activated respiration I_GLU_. The measured HK index value for control cells (12.2%) was very similar to the values measured for the peptide-treated cells **(Table 1)**. The ADP activated respiration was inhibited by the addition of an inhibitor of adenine nucleotide translocator, carboxyatractyloside, which demonstrates the complete intactness of the mitochondrial inner membrane. These data confirm that all four peptides did not affect the Warburg behavior of Caco-2 cells.

**Table 1.** Effects of cell-penetrating FAM-labeled peptides on the coupling of AK- and HK-catalyzed processes with OXPHOS in Caco-2 cells

| Parameters | control | mtCPP1 | UPF25 | mtgCPP | mtCPPgHO |
| --- | --- | --- | --- | --- | --- |
| AK index | 0.50±0.16 | 0.60±0.15 | 0.56±0.19 | 0.53±0.07 | 0.58±0.15 |
| HK index, % | 12.2±1.5 | 13.7±2.4 | 12.2±2.1 | 13.6±1.3 | 11.3±2.0 |

Data are shown as mean values ± SEM, n= 4-5.

**Figure 2.**
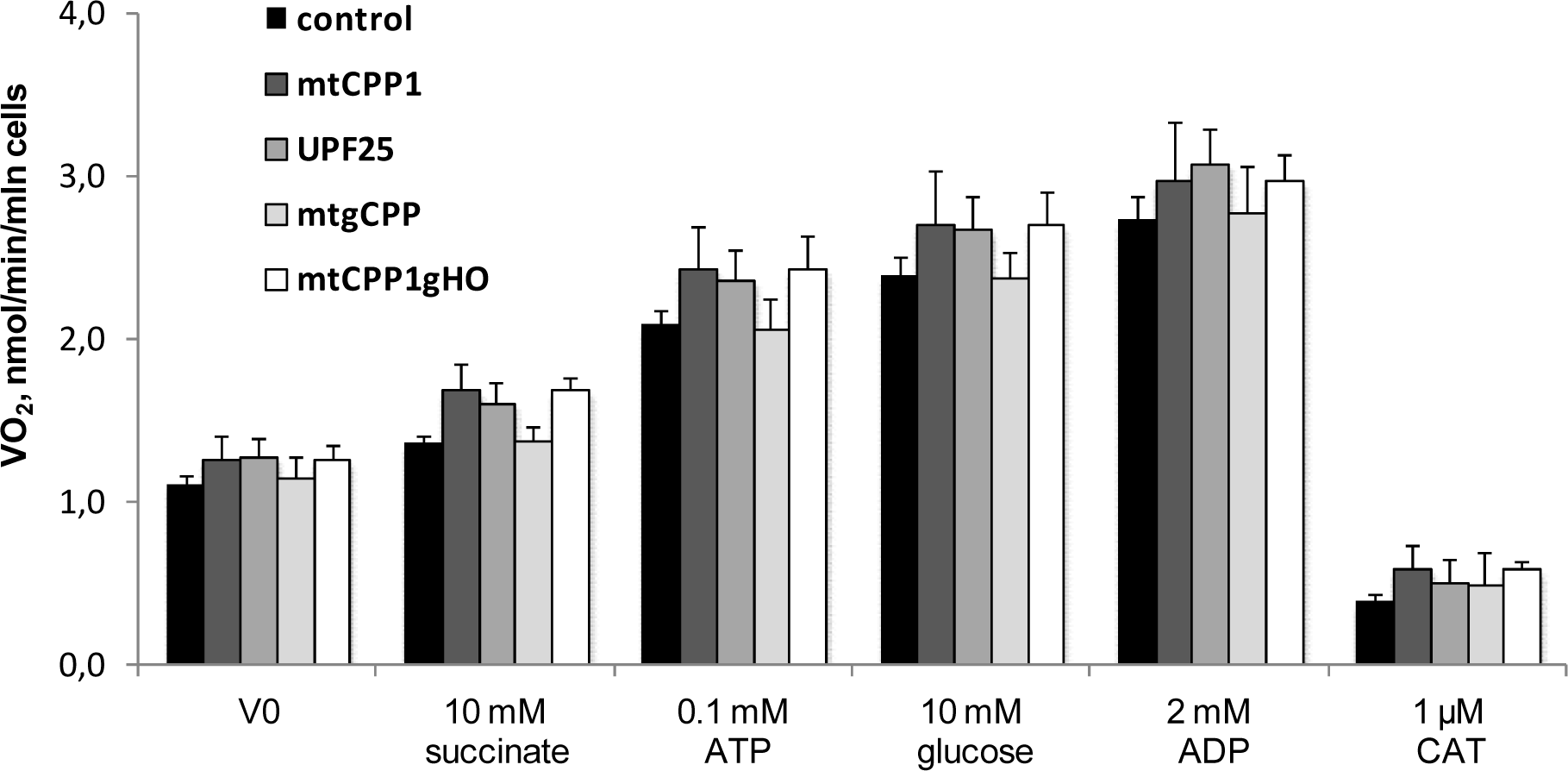
Oxygraphic analysis of the coupling between hexokinase -catalyzed processes and OXPHOS in control and CPPs treated at 5 μM for 24 hrs Caco-2 cells; these experiments were carried out in medium-B with 5 mM glutamate and 2 mM malate as the respiratory substrates. Succinate, glucose, the corresponding adenine nucleotides (ATP or ADP), and carboxyatractyloside (CAT, an inhibitor of adenine nucleotide translocator) were added to the cells sequentially as indicated on the x-axis. VO2 means the rate of oxygen consumption. Data are shown as mean values ± SEM, n = 5.

The AK-catalyzed phosphotransfer system plays a key role in the maintenance of energy homeostasis in well-differentiated cells with intermittent high-energy demand, such as neural, cardiac, and skeletal muscle cells [36]. It is upregulated in human CRC [37] with the total AK activity about 2 times higher than surrounding normal tissue [10]. The same was also observed in tumor cells of other histological types [38]. The AK system could play an important role in the adaptation of cancer cells to an unfavorable microenvironment and tumor progression [39, 40]. Taking into account the role of AK regulated processes in tumor growth we evaluated the effect of CPPs on of this system in Caco-2 cells through its coupling with OXPHOS. Addition of AMP (in the presence of 0.1 mM ATP) to permeabilized cells leads to nearly 30% increase in the rate of O_2_ consumption **(Fig. 3**). Ap5A suppressed the AMP-stimulated respiration, indicating thereby that it was largely mediated by ADP realized in AK reactions. AK index describes the functional activity of AK processes and their coupling with the OXPHOS system. All CPPs did not affect the function of AK system, since the value of I_AK_ measured for control cells (0.50 ± 0.16) did not statistically differ from the value measured for peptide-treated cells (**Table 1)**.

**Figure 3.**
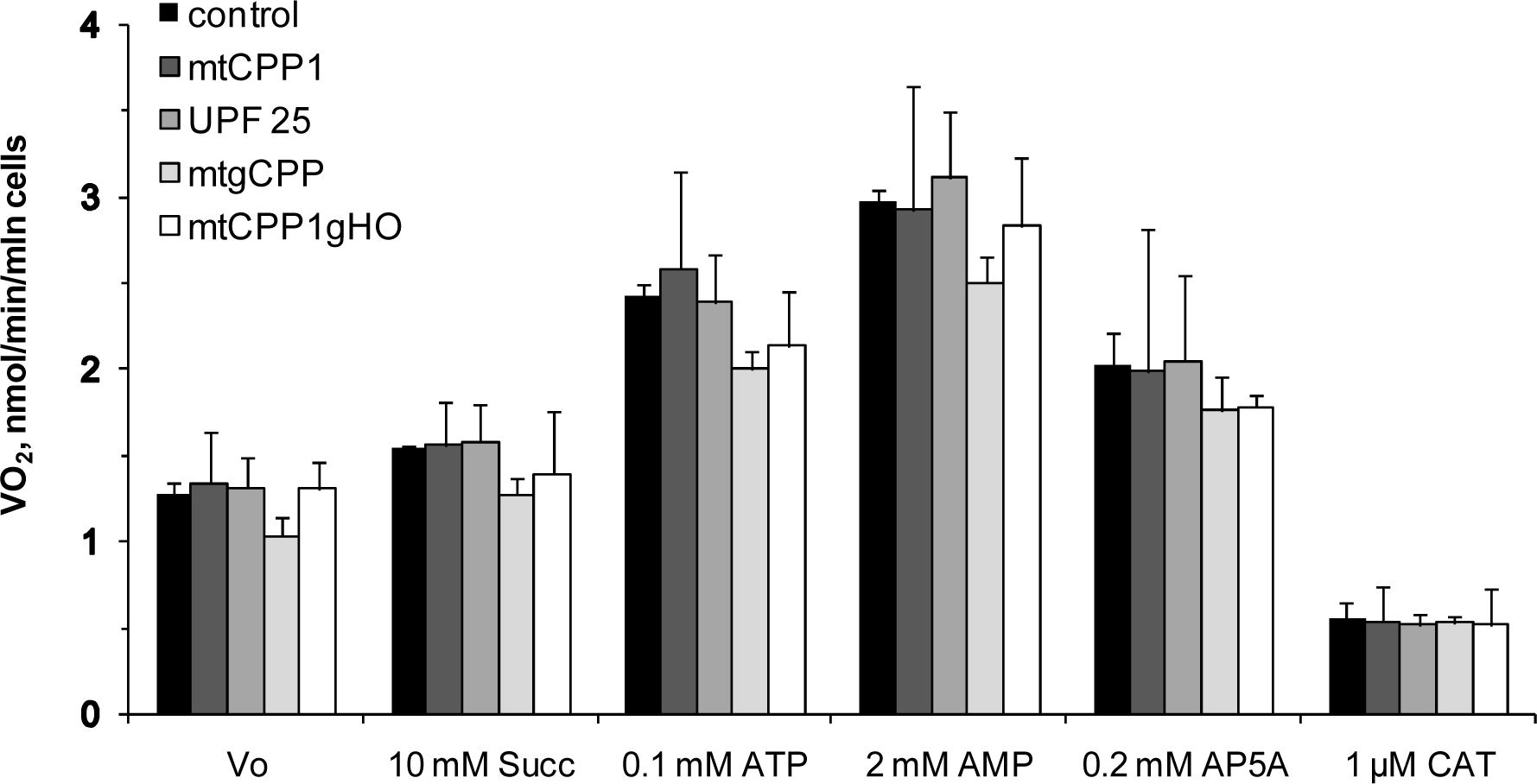
Oxygraphic analysis of the functional coupling between adenylate kinase (AK) catalyzed processes and OXPHOS in control and CPPs pre-treated Caco-2 cells at 5 μM for 24 hrs; addition of 2 mM AMP in the presence of 0.1 mM ATP resulted in activation of mitochondrial respiration due to formation of ADP in AK reactions. The involvement of AK(s) in the stimulation of mitochondrial respiration was confirmed by subsequent addition of diadenosine pentaphosphate (Ap5A), an inhibitor of AK; as can be seen, the addition of Ap5A caused a strong decrease in the rate of O_2_ consumption by these tumor cells. Carboxyatractyloside (CAT) – a selective inhibitor of adenine nucleotide carrier was added to control the intactness of the mitochondrial inner membrane. Data are shown as mean values ± SEM, n = 4-6.

### 3.3. Effect of CPPs on the functional activity of mitochondrial respiratory complexes in Caco-2 cells and the reserve respiratory capacity of these organelles

Previous studies showed that all the studied CPPs can internalize into human cervical carcinoma cells and some other cell lines [23, 31]. Also, mtgCPP and mtCPP1 have the increased ability to target mitochondria without any effect on mitochondrial membrane potential and ATP production by these cells [23]. We proposed that these peptides could affect the bioenergetic function of mitochondria with the elevated OXPHOS activity in CRC cells, unlike glycolytic cervical carcinomas [41, 42] 10, 25, 43].

To investigate the influence of FAM-labeled mtCPP1, UPF25, mtgCPP, and CPP1gHO peptides on the oxygen consumption of the CRC cells and the mitochondrial respiratory system enzymes, we used high-resolution respirometry protocol described previously [44].Mitochondrial respiration in permeabilized Caco-2 cells in the presence of malate and glutamate (LEAK) almost doubled after the addition of 2 mM ADP, indicating the active state of Complex I (**Fig. 4A**). Succinate (final concentration 10 mM) abrogated the strong inhibitory effect of rotenone and exceeded the Complex I maximal rate, showing that the mitochondrial Complex II is more active versus Complex I (**Table 2**). The succinate-fueled respiration was suppressed by the addition of 10 μM antimycin-A, inhibiting the electron flow from Complex III to cytochrome c. Mitochondrial Complex IV was activated by 1.0 mM TMPD, the artificial substrate for reducing cytochrome c in the presence of 5 mM ascorbate, which also shows the total capacity of the respiratory chain (**Fig. 4A**). Finally, the TMPD-activated respiration was inhibited by the addition of 1 mM NaCN, completely inhibiting Complex IV. The mitochondrial respiratory system is the main generator of ROS [45], associated in signaling with cellular stress responses, cell proliferation, and apoptosis [46]. In murine and human tumor models, two different events, mitochondrial respiratory chain overload and partial electron transport inhibition promote superoxide-dependent tumor cell migration, invasion, and metastasis [47, 48]. We found that none of the tested peptides had any effect on the activities of individual respiratory complexes as compared to control cells (Fig.4B). However, in the presence of UPF25 Complex II-linked respiration is significantly higher than mtgCPP (**Fig. 4B**). Besides, a significant increase in the respiratory control ratio (RCR) was detected after treatment with mtgCPP The indicator of the coupling state of mitochondria was calculated as the ratio of maximal ADP-stimulated respiration in the presence of Complex I-linked substrates (glutamate and malate) and the LEAK component. These results indicate a tighter coupling of electron transport and efficient OXPHOS in the CaCo-2 cells treated with the dual antioxidant peptide mtgCPP.

**Table 2.** The influence of FAM-labeled CPPs on the ratio of respiratory Complex I and II in mitochondrial activity

| Parameters of | Caco-2 cells |  |  |  |  |
| --- | --- | --- | --- | --- | --- |
| OXPHOS | control | mtCPP1 | UPF25 | mtgCPP | mtCPP1gHO |
| V<sub>o</sub> | 1.08 ± 0.06 | 1.04 ± 0.06 | 1.18 ± 0.14 | 0.9 ± 0.05 | 1.18 ± 0.07 |
| V<sub>ADP</sub> | 2.07 ± 0.09 | 1.99 ± 0.16 | 2.30 ± 0.19 | 1.94 ± 0.18 | 2.11 ± 0.08 |
| V<sub>Rot</sub> | 0.46 ± 0.02 | 0.38 ± 0.05 | 0.53 ± 0.05 | 0.43 ± 0.06 | 0.47 ± 0.04 |
| V<sub>Suc</sub> | 2.64 ± 0.11 | 2.34 ± 0.29 | 3.05 ± 0.21 | 2.40 ± 0.24 | 2.65 ± 0.16 |
| CI/CII | **0.74** | **0.82** | **0.70** | **0.77** | **0.75** |

**Figure 4.**
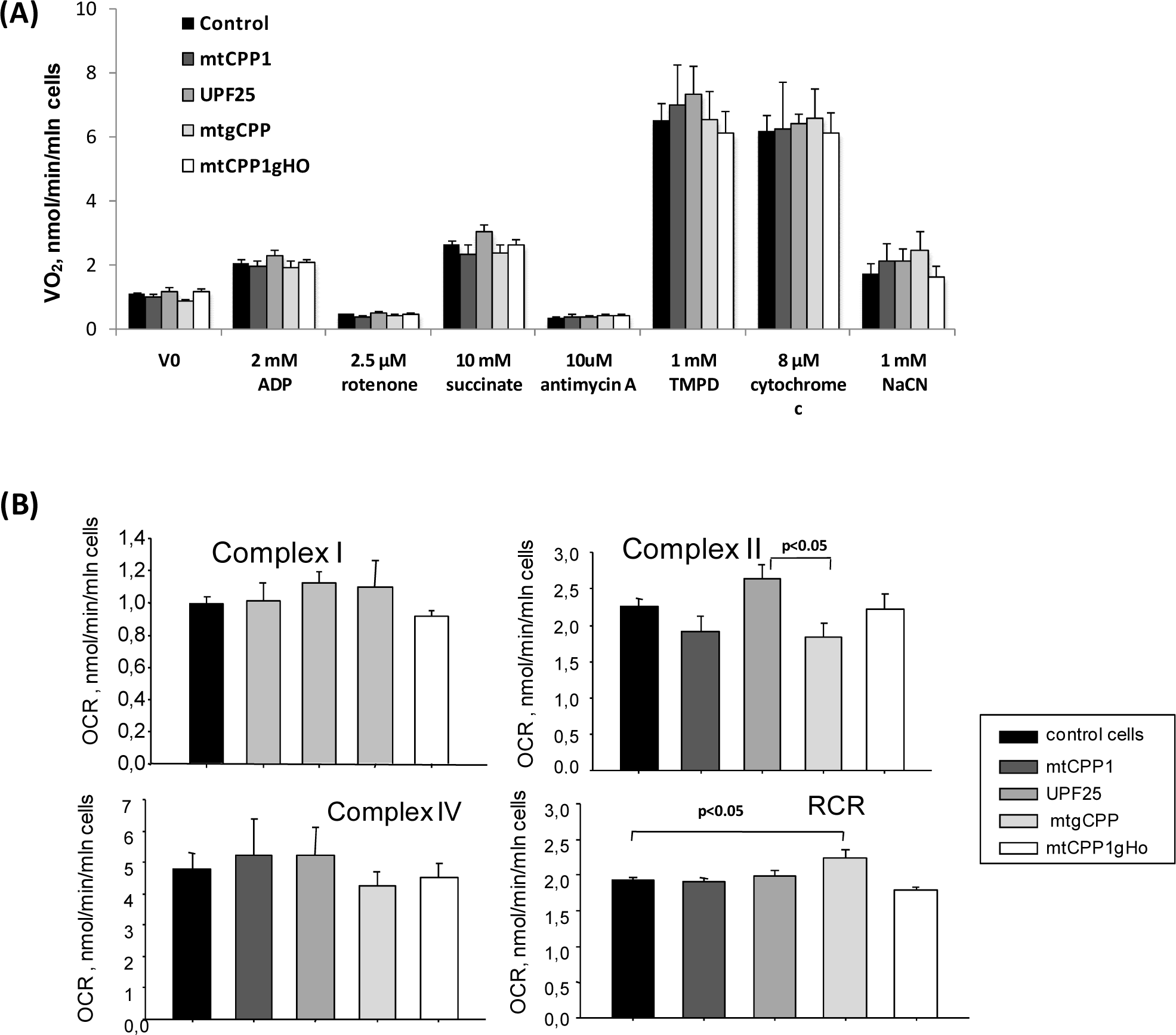
Effect of various CPPs on respiration of the permeabilized Caco-2 cells (**A**) Mitochondrial OXPHOS function test was followed a sequential activation-inhibition of Complexes I-IV of the electron transport system. Oxygen consumption rates expressed as respiration per million cells (nmol/min/mln cells) (**B**) One-way ANOVA indicated no significant differences between mitochondrial respiratory Complexes I, II, IV activities in the CPP-treated and control groups. The respiratory control ratio (RCR) is a measure of mitochondrial coupling state for Complex I. Data are shown as mean values ± SEM, n = 4.

The maximal mitochondrial respiration was measured in the uncoupled state induced by gradual titration with FCCP (p-(trifluoro-methoxy)phenyl-hydrazone) of the untreated and CPP-treated Caco-2 cells according to previously described protocols [49] (**Fig. 5A**). FCCP is a protonophore that uncouples electron transport and mitochondrial respiration from ATP synthesis by disrupting the proton gradient. Respiration rates of control and CPPs treated Caco-2 cells increased about 3 times compared with the proton leak. The optimal concentration of FCCP for peptide treated and untreated cells were 2.21±0.19 μM since at higher concentrations of this uncoupler a decrease in the rate of oxygen consumption was observed. Finally, the electron transport was inhibited by 2.5 μM rotenone and 10 μM antimycin, a Complex III inhibitor, to subtract the non-mitochondrial oxygen consumption. Proton leak and ATP-linked respiration rate are the mitochondrial coupling efficiency indicators. Proton leak is assessed under the condition of the oligomycin - suppressed ATP synthase. None of the peptides tested had any effect of the coupling between ATP synthesis and OXPHOS (**Fig. 5B**). Mitochondrial reserve capacity is the spare ATP amount produced by the mitochondrial respiratory system in case of cell stress or high-energy demand. The summary of the respiratory values had not revealed any alterations (**Fig. 5B**). Statistical analyses confirmed no difference between experimental groups.

**Figure 5.**
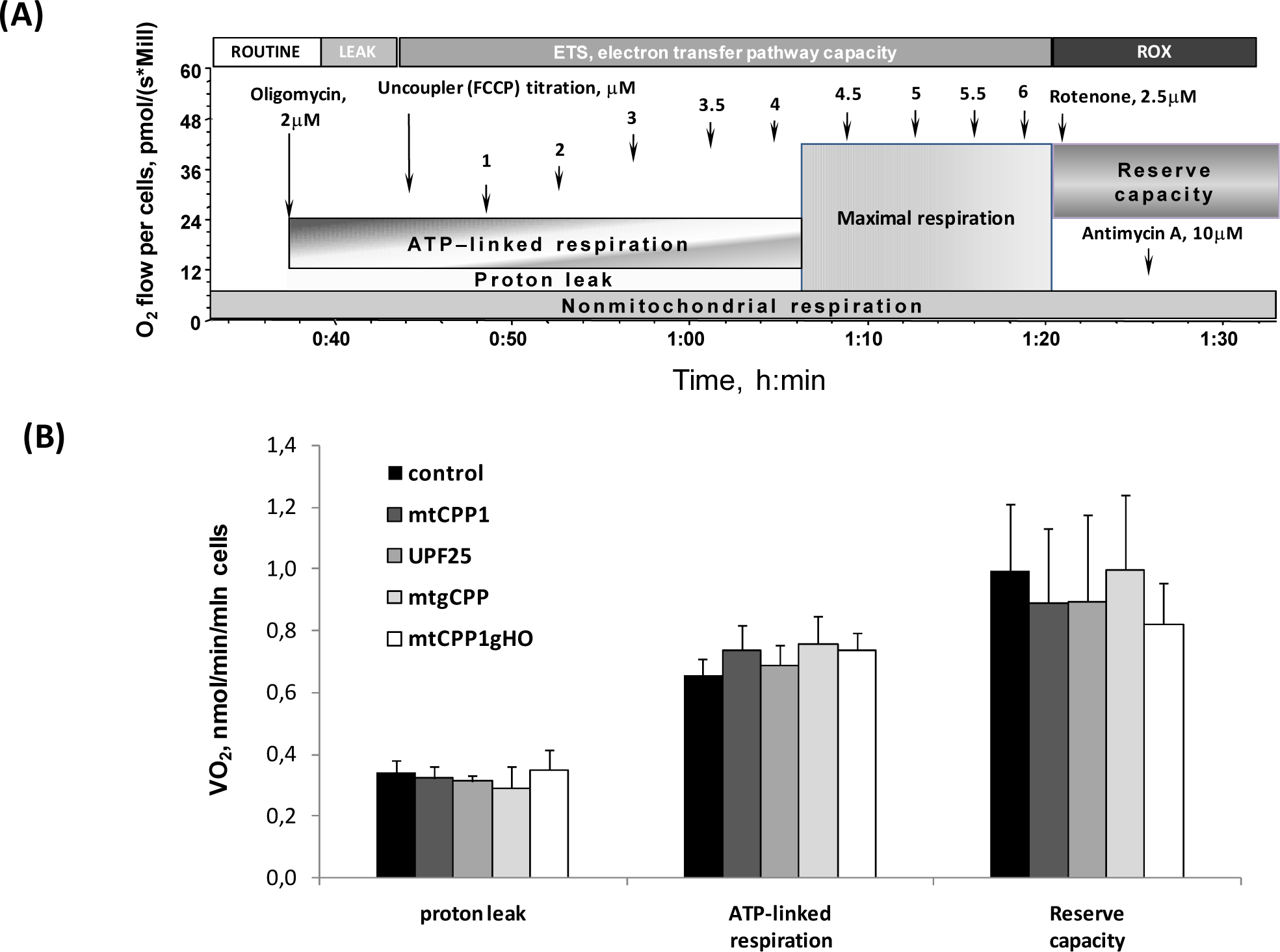
Assessment of mitochondrial function in control and CPP-treated intact Caco-2 cells. (**A**) Representative respirometric traces with a schematic representation of the used mitochondrial respiratory reserve capacity assay; arrows indicate the FCCP concentrations added into the oxygraph chamber. (**C**) Calculated functional properties of mitochondrial respiration in control and CPP-treated Caco-2 cells. These cells were treated with the indicated peptides at a final concentration of 5 μM in complete growth medium for 24 hrs at 37 °C. Data are shown as mean values ± SEM, n = 5.

## 4. Conclusion

None of the tested peptides had any effect on the viability of Caco-2 cells, as well as on their mitochondrial reserve respiratory capacity, the function of respiratory chain complexes, and the small inclination of these cancer cells to aerobic glycolysis. Patients with high levels of mitochondrial markers could be treated with mitochondrial-based therapies in addition to the standard cure to effectively prevent tumor recurrence, metastasis, and drug-resistance [50]. Due to their antioxidative properties and having no negative impact on mitochondrial bioenergetics these mitochondria-targeting CCPs, if conjugated with some specific metabolic regulator for diseased cells, can serve as carrier molecules safe for non-diseased cells. Anti-cancer agents, called “mitocans”, could serve as potential cargo molecules [51] for the studied peptides, which would allow precise delivery of the medication.

In addition to the cancer treatment, these peptides might have potential therapeutic applications in pathologies caused by mitochondrial dysfunctions and aging.

## Abbreviations used

AK: adenylate kinase;
Ap5A: diadenosine pentaphosphate;
CAT: carboxyatractyloside;
CRC: colorectal cancer;
CPPs: cell-penetrating peptides;
HK: hexokinase;
FAM: 5-(6)-carboxyfluorescein;
MTT: 3-(4,5-dimethylthiazolyl-2)-2, 5-diphenyltetrazoliumbromide;
OXPHOS: oxidative phosphorylation;
ROS: reactive oxygen species;
TMPD: N,N,N’,N’-tetramethyl-phenylenediamine;
VDAC: voltage dependent anion channel.

